# Independent dynamics of slow, intermediate, and fast intracranial EEG spectral activities during human memory formation

**DOI:** 10.1101/2021.05.06.442655

**Authors:** Victoria S. Marks, Krishnakant V. Saboo, Çağdaş Topçu, Theodore P. Thayib, Petr Nejedly, Vaclav Kremen, Gregory A. Worrell, Michal T. Kucewicz

**Affiliations:** Graduate School of Biomedical Sciences, Mayo Clinic; University of Illinois, Dept. of Electrical and Computer Engineering, Urbana-Champaign IL, USA; Gdansk University of Technology, Faculty of Electronics, Telecommunications and Informatics, BioTechMed Center, Multimedia Systems Department, Gdansk, Poland; Nencki Institute of Experimental Biology, Polish Academy of Sciences, Warsaw, Poland; Mayo Clinic, Dept. of Neurology, Rochester MN, USA; Iowa State University, Department of Computer Engineering, Ames, Iowa, USA; The Czech Academy of Sciences, Institute of Scientific Instruments, Brno, Czech Republic; Czech Institute of Informatics, Robotics, and Cybernetics, Czech Technical University in Prague, Prague, Czech Republic; Department of Physiology and Biomedical Engineering, Mayo Clinic

## Abstract

A wide spectrum of brain rhythms are engaged throughout the human cortex in cognitive functions. How the rhythms of various low and high frequencies are spatiotemporally coordinated across the human brain during memory processing is inconclusive. They can either be coordinated together across a wide range of the frequency spectrum or induced in specific bands. We used a large dataset of human intracranial electroencephalography (iEEG) to parse the spatiotemporal dynamics of spectral activities induced during formation of verbal memories. Encoding of words for subsequent free recall activated slow theta, intermediate alpha and beta, and fast gamma frequency power in discrete cortical sites. A majority of the electrode sites recorded activity in only one of these frequencies, except for the visual cortex where spectral power was induced across multiple bands. Each frequency band showed characteristic dynamics of the induced power specific to cortical area and hemisphere. The power of the low, intermediate, and fast activities propagated in distinct spatiotemporal patterns across the visual, temporal and prefrontal cortical areas as the words were presented for encoding. Our results suggest anatomically and temporally distributed spectral activities in the formation of human memory.

## 1. Introduction

Brain rhythms are thought to support memory and cognitive functions by coordinating activities of connected neurons into synchronous interactions (Singer 1999; Buzsaki 2006; Fries 2015; Klimesch 1996; Fell and Axmacher 2011). The spectrum of rhythmic activities involved in these interactions includes the theta (3-9 Hz) and the gamma (30-120 Hz) frequency bands with specific roles proposed for the slow and fast activities in memory processing (Düzel, Penny, and Burgess 2010; Sauseng et al. 2010; Fries, Nikolić, and Singer 2007; Tallon-Baudry and Bertrand 1999; Nyhus and Curran 2010; Lisman and Jensen 2013). The intermediate alpha (9 - 12 Hz) and beta (12 - 25 Hz) frequency bands were also associated with processes necessary for memory functions (Klimesch, Sauseng, and Hanslmayr 2007; Hanslmayr et al. 2011; Spitzer and Haegens 2017; Schmidt et al. 2019; Michalareas et al. 2016). How these various rhythms are operating together in the functional anatomical space of specific brain regions and during the temporal phases of memory processing remains poorly understood. The synthesis of these previous studies provides two possible scenarios. In one scenario, the entire spectrum of brain electrophysiological activities is engaged in particular areas responsible for processing information at a whole range of frequency timescales. In another scenario, each area is responsible for processing information at a defined timescale, engaging only a subset of activities in the frequency spectrum. Both scenarios agree that the brain activities at defined frequency ranges can be conceptualized as “spectral fingerprints” that subserve specific perceptual or cognitive processes at a corresponding temporal scale of neural interactions (Siegel, Donner, and Engel 2012; Hanslmayr and Staudigl 2014). Tracking the dynamics of these spectral fingerprints accurately in the anatomical space and time is critical to test these hypotheses and shed light on the local and global processes engaged during memory tasks.

Intracranial electroencephalographic (iEEG) recordings present a unique opportunity to probe with high resolution the large scale of neural activities engaged during cognitive and other brain functions in discrete areas of the human cortex (Lhatoo, Kahane, and Luders 2019; Johnson et al. 2020; Jacobs and Kahana 2010; Engel et al. 2005). The iEEG signals are sampled from electrode contacts implanted either directly on the surface of the neocortex or into its deeper layers and subcortical structures. Thus, spectral activities generated by local neural populations and recorded from multiple separate contacts can be tracked in discrete cortical areas during the time of memory processing. Previous iEEG studies employed predominantly the spectral power induced in high gamma frequency ranges (60-120 Hz) to track cortical processing in memory and cognitive tasks (Jerbi et al. 2009; Lachaux et al. 2012; Crone, Sinai, and Korzeniewska 2006; Lundqvist et al. 2018). Other task-induced iEEG activities in the lower frequency bands were less commonly investigated. Oscillatory activities in the theta and alpha frequencies were shown to propagate as local, independent, travelling waves in discrete cortical areas during memory processing (Zhang et al. 2018) but the relative spatiotemporal dynamics of these slow and the faster beta and gamma activities during memory processing remains largely unexplored.

Here, we took advantage of a large iEEG dataset of 164 participants encoding lists of words for subsequent free verbal recall. Previous studies with this task revealed a distributed network of brain regions associated with sensory processing and with higher-order declarative memory functions (J. F. Burke et al. 2013; John F. Burke, Long, et al. 2014; Kucewicz et al. 2019). The induced high gamma power was observed first in the visual areas of the occipito-temporal cortex and then in more anterior areas of the prefrontal cortex, inspiring a two-stage model of memory encoding with an early sensory and a late association stage (John F. Burke, Long, et al. 2014). A similar sequence of posterior-to-anterior order of induced power was also observed in the theta frequency band (J. F. Burke et al. 2013), and also confirmed in activities beyond the gamma frequency range (Kucewicz et al. 2014). Our recent study found the sequence of the induced high gamma power during memory encoding to be continuous with increasing latencies along a hierarchy of gradually higher-order association areas (Kucewicz et al. 2019), culminating in the anterior prefrontal cortex late into word encoding. The previous studies were either limited to smaller datasets of participant recordings with sparse electrode coverage of the neocortex or focused only on selected frequency bands. Hence, spectral power across various bands would either be averaged spatially across multiple electrode sites of larger cortical regions, thereby losing the resolution of individual sites, or would not be studied across the slow theta, intermediate alpha and beta, and fast gamma brain activities.

Given the wide scope of the frequency spectrum and brain areas analyzed collectively in the previous studies, we hypothesized that specific spectral activities are induced in a pattern of discrete cortical locations. Hence, aggregating activity from several locations gives rise to a broadband tilt in power across the entire spectrum (Kilner et al. 2005), which is actually composed of multiple specific ‘spectral fingerprints’ (Fellner et al. 2019). Exact localization of different fingerprints would either be common to the same cortical site of electrode recording or would be distributed in a mosaic of distinct anatomical sites. Likewise, in terms of the timing, the slow, intermediate and fast frequency spectral activities would be induced at different times of memory encoding or all at the same time. In addition to the theta and gamma activities, alpha and beta rhythms were expected to be induced at distinct cortical sites and times of memory encoding. In general, we tested for independent spatiotemporal dynamics of the spectral activities induced during formation of human verbal memory traces.

## 2. Materials and Methods

### 2.1. Subjects

The dataset of iEEG recordings from a total of 164 participants undergoing surgical evaluation of drug-resistant epilepsy was taken from a large multicenter collaborative study (all de-identified data are available at http://memory.psych.upenn.edu/Electrophysiological_Data). The recordings were collected from the following centers: Mayo Clinic, Thomas Jefferson University Hospital, Hospital of the University of Pennsylvania, Dartmouth-Hitchcock Medical Center, Emory University Hospital, University of Texas Southwestern Medical Center, and Columbia University Hospital. One common research protocol was approved by the respective Institutional Review Board at each center, and informed consent was obtained from each participant. Electrophysiological recordings were collected from standard clinical subdural and depth electrodes (AdTech Inc., PMT Inc.) implanted on the cortical surface and into the brain parenchyma respectively. Subdural electrode contacts were arranged either in a grid or a strip configuration with 10 mm separation, and depth electrode contacts were separated by 5 to 10 mm. The placement of electrodes was determined by a clinical team with the goal of localizing seizure foci for possible epilepsy resective surgery or implantation of a device for therapeutic electrical brain stimulation.

### 2.2. Anatomic Localization and Brain Surface Mapping

Cortical surface parcellations were generated for each participant from pre-implant MRI scans (volumetric T1-weighted sequences) using Freesurfer software (RRID:SCR_001847). The hippocampus and surrounding cortical regions were delineated separately based on an additional 2 mm thick coronal T2-weighted scan using the Automatic Segmentation of Hippocampal Subfields (ASHS) multi-atlas segmentation method. Electrode contact coordinates derived from co-registered post-implant CT scans were then mapped to the pre-implant MRI scans to determine their anatomic locations. For subdural strips and grids, the electrode contacts were additionally projected to the cortical surface using an energy minimization algorithm to account for postoperative brain shift. Contact locations were reviewed and confirmed on surfaces and cross-sectional images by a neuroradiologist. The T1-weighted MRI scans were also registered to the MNI152 standard brain to enable comparison of recording sites in a common space across subjects. Anatomic locations of the recording sites, including Brodmann areas, were derived by converting MNI coordinates to Talairach space and querying the Talairach daemon (www.talairach.org).

### 2.3. Electrophysiological Recordings

The iEEG signals were recorded using one of the following clinical electrophysiological acquisition systems (dependent on the institution for data collection): Nihon Kohden EEG-1200, Natus XLTek EMU 128, or Grass Aura-LTM64. Depending on the acquisition system and the preference of the clinical team, the signals were sampled at either 500, 1000, or 1600 Hz and were referenced to a common contact placed either intracranially, on the scalp, or on the mastoid process. For analysis, all recordings using higher sampling rates were filtered by an antialiasing filter and down-sampled to 500 Hz. A common bipolar montage was calculated *post hoc* for each subject by subtracting measured voltage time series on all pairs of spatially adjacent contacts. This resulted in N-1 bipolar signals in case of the penetrating and the strip electrodes, and N = (i-1)*j + (j-1)*i, where i and j are the numbers of contacts in the vertical and horizontal dimensions of the grid. For the data analysis of this study, one “electrode” refers to the bipolar signal from one bipolar pair of contacts. It is important to note that contacts may be used for multiple electrodes, so not all signals are truly independent.

### 2.4. Free Recall Task

A classic paradigm for probing formation of verbal episodic memory was employed (Kahana 2014), in which subjects were shown lists of words for subsequent free recall. Participants were instructed to study lists of words presented on a laptop computer screen for a delayed test of vocal recall. Lists were composed of 12 words chosen at random and without replacement from a pool of high-frequency nouns in the subject’s native language (either English or Spanish; http://memory.psych.upenn.edu/WordPools). Each word appeared on the screen for 1600 ms, followed by a random jitter of 750 to 1000 ms blank interval between stimuli. At the end of each word list, a distractor task was performed by the subject. This task lasted for 20 s and was composed of a series of simple, arithmetic problems of the format A + B + C, where A, B, and C were random, single-digit integers between 1 and 9. After the distractor task, participants were given 30 s in which to recall as many of the words from the list as possible in any order (Figure 1). Vocal responses were recorded digitally by the laptop and were later scored manually for analysis. Only subjects who recalled at least 15% of words and completed at least 12 lists of the task were included in further analysis. This left 139 of the 164 original subjects for a grand total of 14,219 electrodes used in this study. Electrophysiological recordings were synchronized to stimulus appearance on the screen through an electric pulse generator operated by the task laptop, which sent pulses to a designated event channel in the clinical acquisition system. The events were timestamped after the recording session using custom-written MATLAB codes and were used to extract specific epochs of interest around word presentation. Each recorded epoch was 3000 ms long and included 1600 ms of word presentation on the screen with 700 ms of blank screen of interstimulus interval before and after each word presentation.

**Figure 1.**
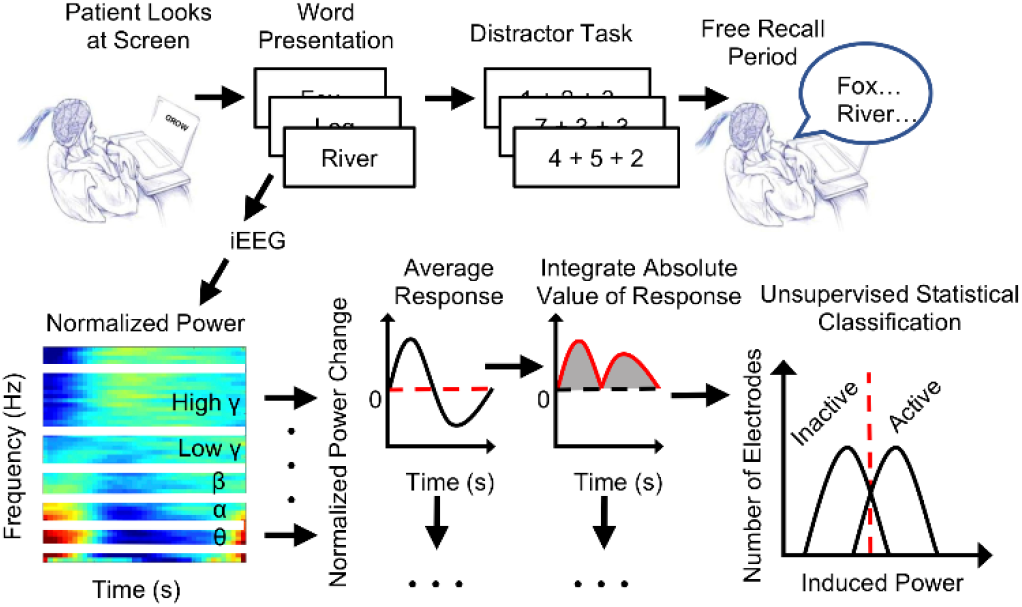
Fully-automated classification of induced spectral power identifies active electrodes during verbal memory encoding. Trial-averaged spectral power of iEEG signals from epochs of word presentation (one word per trial) was integrated across the epoch and used as a feature to classify active electrodes that record from brain regions engaged in verbal memory encoding. Active electrodes were classified independently in six frequency bands of the iEEG spectrum using normalized estimates of the power change (z-score transform). The classification procedure used an unsupervised method based on the Gaussian Mixture Model (Saboo et al. 2019). Notice induced activity across all six frequency bands in the example spectrogram plotted from trial-averaged activity from one electrode localized in the occipital cortex.

### 2.5. Data Analysis

We analyzed the iEEG recording epochs during word presentation for memory encoding. Each presentation epoch was FIR filtered (2000-order Barlett-Hanning with zero-phase distortion filtering, bandpass with frequency limits specified for each frequency band) before being spectrally decomposed, normalized, and binned independently into distinct frequency bands between 2 and 120 Hz: low theta (2-4 Hz), high theta (5-9 Hz), alpha (10-15 Hz), beta (16-25 Hz), low gamma (25-55 Hz), and high gamma (65-115 Hz). Spectrograms of the word presentation epoch were averaged together after log-normalization and a z-score transformation of each time-frequency point. Z-scoring was performed according to the following formula:

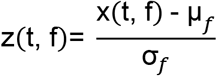

where x is the signal, t is the time bin, f is the frequency bin, μ_f_ is the mean, and σ_f_ is the standard deviation for the given frequency. Each trial was normalized separately. Log normalization was implemented in order to correct for the 1/f-power law effect of lower frequencies on estimating power in the higher frequency bands, and the z-score transformation was performed to provide a normalized scale of power change for comparison of signals from different electrodes, sessions and participants (Kucewicz et al. 2019; Saboo et al. 2019; Kucewicz et al. 2017). This z-score normalization produces positive and negative power changes relative to the mean power within any one word epoch instead of using a “baseline” period. Hence, even small positive or negative deviations of absolute power would effectively be augmented relative to the signal mean, as described before (Kucewicz et al. 2019; Alotaiby et al. 2015). Edge artifacts in the power estimate were eliminated by clipping two time bins from either end of the final spectrogram for all frequencies. For each electrode and frequency band, we determined the average spectral power at each time bin across all word presentation epochs. The overall induced power in a given frequency band during the word presentation period was quantified as the area under the absolute value of the corresponding time-series curve, as shown in Fig. 1. Electrodes were then classified into “active” and “inactive” in each specific band using our unsupervised clustering method based on Gaussian Mixture Modeling (GMM) to identify those that showed task-induced power changes (Saboo et al. 2019). We chose a GMM-based method to ensure that different numbers of electrodes would be allowed for the clusters of active electrodes and the inactive electrodes. The number of clusters was set to two because the active-inactive categorization of electrodes is binary: change in the spectral power can either be induced or not during the word presentation period. The use of a GMM also enabled classification without a requirement for ground truth data. Electrodes were pooled across subjects and then clustered to identify active electrodes.

### 2.6. Induced Power Mapping

Most of the active electrodes were localized in the cortical regions associated with processing visual and semantic information, as well as with declarative memory and executive functions. Out of 38 Brodmann Areas, the majority of electrode locations grouped into 9 brain regions each comprising Brodmann Areas: visual (Brodmann areas 18 and 19), inferior temporal (Brodmann areas 20 and 37), precuneus (Brodmann areas 30 and 31), lateral parietal (Brodmann areas 7 and 39), mesial temporal (Brodmann area 28 and Hippocampus), lateral temporal (Brodmann areas 21 and 22), Broca’s areas (Brodmann areas 44 and 45), lateral prefrontal (Brodmann areas 9 and 46), and frontal pole (Brodmann areas 10 and 11). Unless stated otherwise, all further analysis was focused on the recalled word epochs, i.e. epochs with successfully encoded words that were subsequently freely recalled, in these nine cortical regions.

Induced power changes in each frequency band were plotted from all active electrodes onto an average brain surface for each time point of the word epoch using a custom MATLAB code for an electrode-location-dependent weighing technique. The brain surface was created as a mesh, where each element of the mesh was assigned a color dependent on the induced power of the surrounding electrodes weighed by the relative distance to the element (greater weight to more proximal electrodes). For each mesh element of the brain surface plotted, only the effects of electrodes within a threshold of 10 millimeters from the element center point were considered. Hence, the induced power was averaged from all electrodes within the threshold radius that were linearly weighted by their proximity according to the following formula:

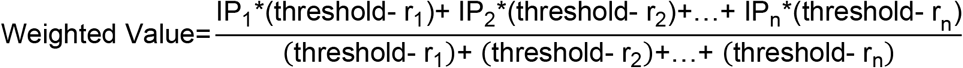

where IP is the induced power change value for an electrode, and r is the distance between the electrode and the chosen element of the brain surface mesh.

### 2.7. Statistics

ANOVA and multiple comparison tests were used to analyze the effects of frequency band, brain region (Brodmann Area), and hemisphere on the percentage of electrodes in an area labeled as “active.” Temporal differences between hemispheres were quantified using a two-tailed, unpaired t-test statistic with Bonferroni correction. We then broke each word epoch into pre-encoding, early encoding, late-encoding, and post-encoding phases and used repeated measures ANOVA with post-hoc Tukey-Kramer test to analyze the effects of frequency band, brain region, and hemisphere on induced power during each phase of the verbal memory task. Peak latency for each frequency and area was determined as the time bin with the maximum positive peak of the induced power. Latencies were compared using repeated measures ANOVA and post-hoc Tukey-Kramer multiple comparison tests. The alpha value for all tests was set at 0.05.

## 3. Results

### 3.1. Spectral activities induced during memory encoding are heterogeneously distributed across the human cortex

We first identified a subset of electrodes from cortical areas activated during memory encoding. For this purpose, spectral power-in-band in six non-overlapping frequency ranges (low theta: 2-4 Hz, high theta: 5-9 Hz, alpha: 10-15 Hz, beta: 16-25 Hz, low gamma: 25-55 Hz, high gamma: 65-115 Hz) was used as features for automated classification of active electrodes (Saboo et al. 2019) independently in each band (Fig. 1). Out of the total of 14,219 electrodes implanted across all participants, 4738 (33.3%) showed the induced spectral power in at least one of the frequency bands. In agreement with our previous results (Kucewicz et al. 2019; Alotaiby et al. 2015), these active electrodes were widely distributed across 39 Brodmann areas that were defined in all cortical lobes (Fig. 2A). The relative proportion of electrodes in each Brodmann area that were active varied from 4.3% to 75%, depending on location within the limbic, frontal, prefrontal, parietal, temporal, or occipital lobes (ANOVA F=99.21, p < 0.001, df = 5), with significantly more active electrodes found in the sensory visual areas of the occipital and the parietal cortex (post-hoc Tukey Kramer, p<0.05). Within each area, similar proportions were observed in each frequency band, including the alpha and the beta band, with no significant differences across the spectrum (N-way ANOVA F=0.17, p = 0.9747, df = 5). In summary, regardless of frequency band analyzed, at least 4% of electrode sites within any of the cortical areas analyzed revealed significantly induced power during the task.

**Figure 2.**
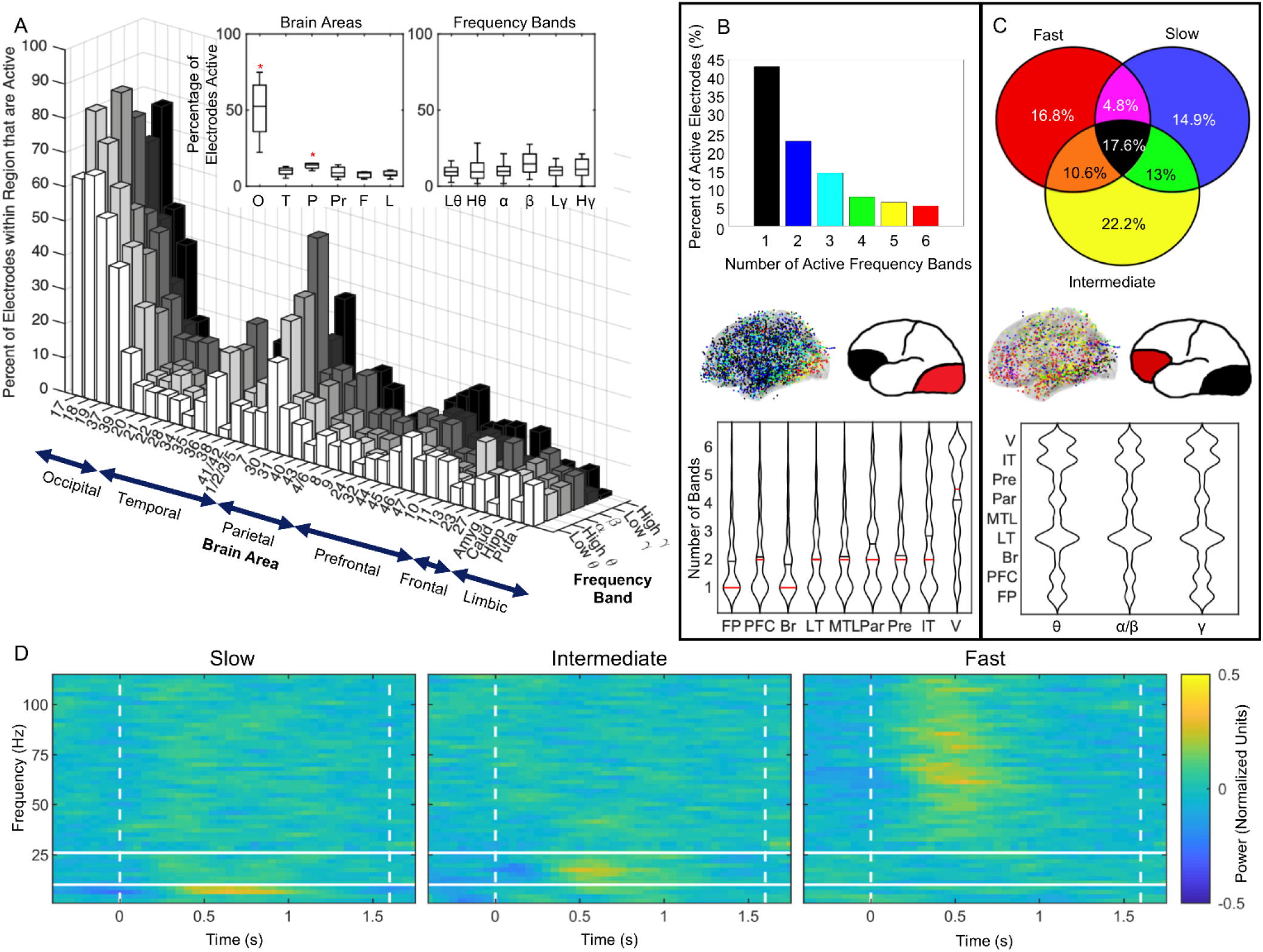
Spectral activities induced during verbal memory encoding are non-uniformly distributed across the human brain. (A) Proportions of active electrodes in six frequency bands studied (low theta: 2-4 Hz, high theta: 5-9 Hz, alpha: 10-15 Hz) beta: 16-25 Hz, low gamma: 25-55 Hz, high gamma: 65-115 Hz) show significant differences in the anatomical distribution (red asterisks; p<0.05 post-hoc Tukey Kramer test) but not across the frequency bands as summarized in the inset box plots. (B) Distribution of the electrodes active in one or more of the frequency bands reveals the highest overlap in the posterior areas of the occipital and parietal visual cortex, which was gradually decreasing in the more anterior cortical areas as shown in the average brain surface plot and the cartoon summary of all active electrodes (dots colored according the bar chart) and in the violin plots (black and red lines indicate the mean and median, resp. (Hoffmann, 2018)). (C) Distribution of all electrodes active in the gamma (fast), alpha/beta (intermediate), and theta bands (slow) differs across nine selected cortical regions of interest (V - visual; IT - inferior temporal; Pre - precuneus; Par - lateral parietal; MTL - mesial temporal lobe; LT - lateral temporal; Br - Broca’s area; PFC - prefrontal cortex; FP - frontal pole) with relatively more prefrontal electrodes active in the gamma bands (red) and more electrodes in the visual areas active across all the bands (black) as shown in the average brain surface plot, the cartoon summary, and the violin plot. Notice that most electrodes overall were active in just one or two frequency bands either of the slow, intermediate or fast frequencies, except for the visual areas with more than two frequency bands active per any one electrode. (D) Representative examples of trial-averaged spectrograms during word encoding from active electrodes show the induced power changes confined to each of the three frequency groups from different anatomical locations.

To investigate whether these various slow and fast spectral activities were induced on the same electrodes, we summarized the overlap across the six frequency bands for nine cortical regions of interest (ROI; each composed of two Brodmann areas with >18 active electrodes from >8 participants; see Table 1). An electrode can be active in one or more bands. We found that the highest proportion of electrodes were active in only one (43.12%) or in two (22.94%) bands, and proportionally fewer in more than two bands with only a minority (5.4%) active in all six (Fig. 2B). The highest overlap of induced spectral power in more than two bands was observed for the electrodes in the posterior sensory visual areas of the occipital cortex Fig. 2B; red), where 4-5 bands were activated on average per electrode during word encoding. Other visual areas of the temporal and parietal cortex also showed relatively high distributions of electrodes with more than two bands activated, in contrast to the anterior higher order processing areas in the temporal and prefrontal cortex, where electrodes were active in 1-2 frequency bands (Fig 2B; black). Overall, the majority of electrode sites showed the induced spectral power in one or two specific frequency bands, suggesting that the various activities were spatially segregated in the cortex.

**Table 1.**
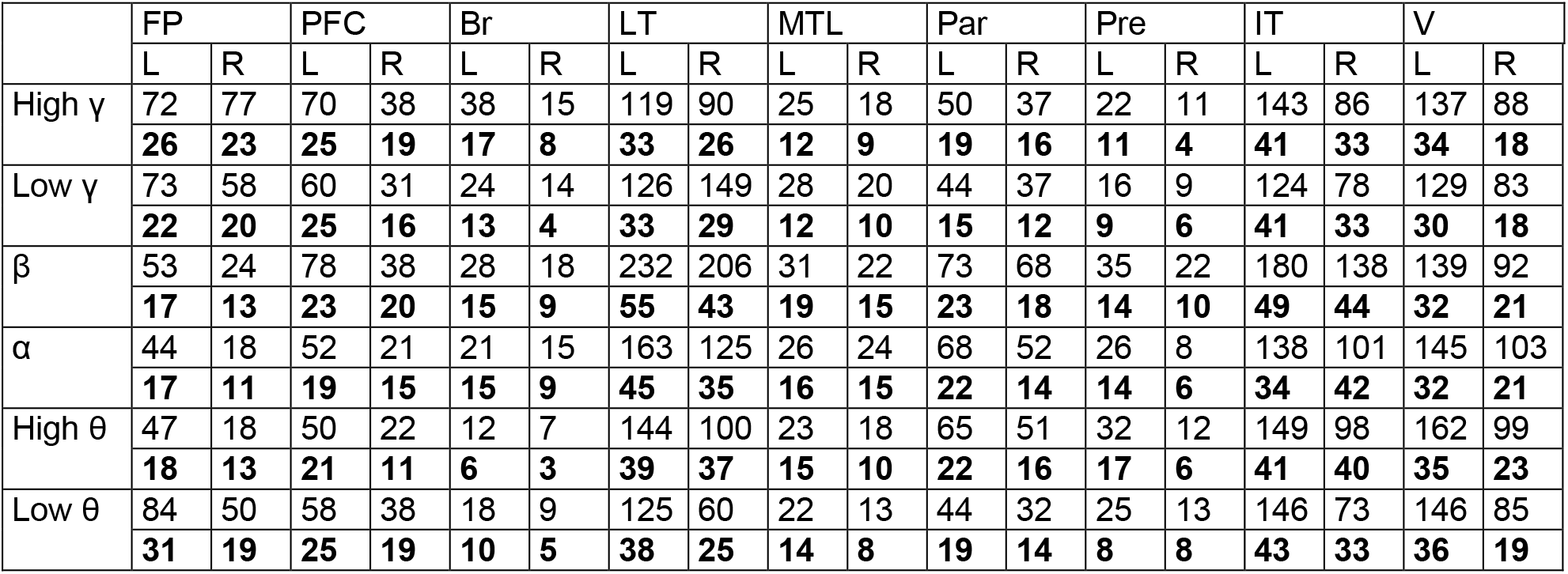
Summary of the total number of active electrodes used in the analysis. Electrodes separated by location (including hemisphere) and frequency band of activity. The number of participants that these came from are included in bold. (FP – Frontal Pole, PFC – Prefrontal Cortex, Br – Broca’s Area, LT – Lateral Temporal, MTL – Mesial Temporal Lobe, Par – Lateral Parietal, Pre – Precuneus, IT – Inferior Temporal, V – Visual, L – Left Hemisphere, R – Right Hemisphere).

We next asked whether the spectral activities that were induced at any one electrode site overlapped in similar frequency ranges. We grouped the activities into slow, intermediate, and fast frequency groups (theta, alpha/beta, and gamma, respectively) and compared proportions of active electrodes with different group combinations. More than half of all active electrodes showed induced power in exclusively the slow, the intermediate, or the fast frequencies (Fig. 2C). Electrodes with high overlap across all groups were predominantly localized in the posterior visual areas (Fig. 2C; black). The remaining electrode sites that showed induced power in one of the three frequency groups were uniformly distributed across the cortex with similar proportions in each ROI, apart from the prefrontal areas where relatively more fast gamma activities were present during memory encoding (Fig. 2C; red). Our results showed that activities of particular frequency range were induced mostly at specific cortical sites, forming a mosaic-like pattern of distribution consistent with the notion of ‘spectral fingerprints’ (Siegel, Donner, and Engel 2012). Hence, a typical electrode site showed induced power in a defined range of either slow, intermediate or fast frequencies (Fig. 2D).

### 3.2. Laterality effect of the induced power is specific to particular frequencies and cortical areas

Our next question was if the total power-in-band induced during the time of word encoding (Fig. 1) was different depending on the anatomical location, the frequency band, or the hemisphere. Individual active electrodes can show similar magnitudes of the induced power, despite the heterogeneous distribution across the cortical regions (Fig. 2). Overall, there was a significant effect of the frequency band on the induced power (repeated measures ANOVA F=1346.99, p<0.001, df=5). Brain region (ANOVA F=38.21, p<0.001, df=37) and of the hemisphere location of the electrode (ANOVA F=17.4, p<0.001, df=1). Post-hoc group analysis (Tukey-Kramer, p<0.05) confirmed significantly smaller power with increasing frequencies of the bands, except between the low and high gamma bands (Supplementary Fig. 1). Electrodes within the occipital lobe showed significantly higher induced power than in any other area in agreement with our previous study of the high gamma activities (Kucewicz et al. 2019).

In general, the total induced power was significantly different between the left and the right hemisphere. We investigated this laterality effect by estimating the temporal profiles of the mean power change in each ROI. While there was a significant laterality effect in at least one frequency band of every ROI (Fig. 3A; t-statistic, p<0.05), the timing and magnitude of this effect was specific to a particular band and ROI. The temporal profiles of the induced power in the left and the right hemisphere (Fig. 3A; red and blue, resp.) provide another spectral signature of particular cortical areas. Hemisphere had an effect on the induced power in time (Fig. 3B; repeated measures ANOVA, encoding phase as within subjects factor, F=17.01, p<0.001, df = 3). When the total induced power was compared between the brain regions (post-hoc Tukey-Kramer, p<0.05) only the prefrontal, inferior temporal, and visual cortex showed a significant laterality effect (Fig. 2B). The effect in these three areas was significant regardless of the frequency band analyzed. Significantly more power in the left prefrontal and inferior temporal areas was expected given the role of these areas in processing verbal information.

**Figure 3.**
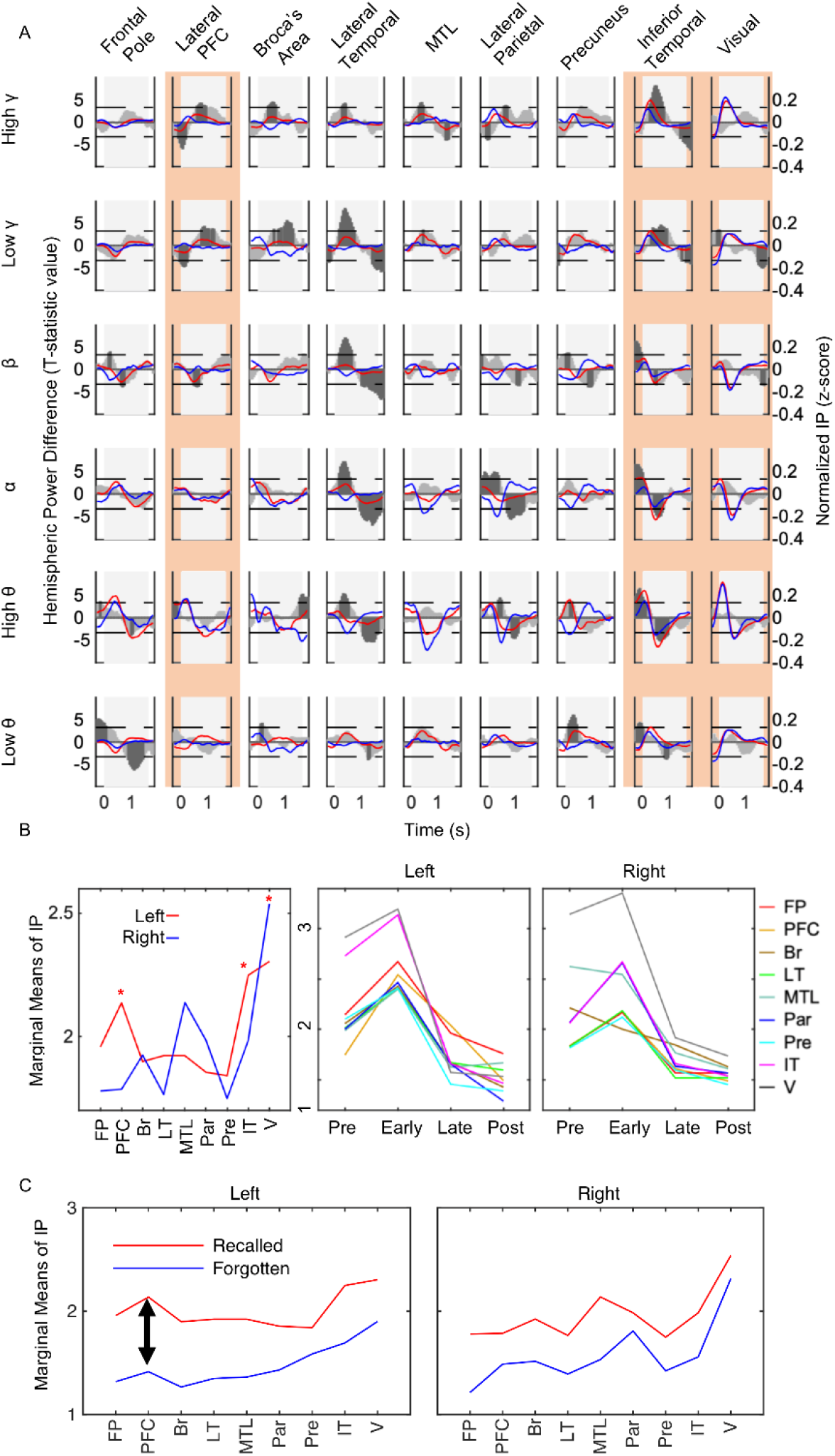
Temporal profiles of induced power (IP) during memory encoding reveal a strong laterality effect in specific frequency bands and cortical areas. (A) Each panel shows trial-averaged power changes across the time of word presentation (gray background) estimated from all active electrodes in a given frequency band and brain region plotted separately for the left and right hemisphere (red and blue, resp.). The gray area plots in the background summarize the statistical difference (t-statistic) between the two hemispheres in 50 ms time bins with dark gray indicating significantly more (positive t values) or less (negative t values) power in the left hemisphere (horizontal dashed lines; Student t-test, p<0.05). Notice that the significant laterality effect localized in specific frequency bands in a given brain region. (B) Summary comparison of the total induced power from all frequency bands across the nine brain regions (see Fig. 2 for labels) and four stages of memory encoding (red asterisks indicate significant difference; Tukey-Kramer test, p<0.05). (C) Memory effect was the largest in the left prefrontal cortex (indicated by black double arrow), whereas, MTL showed the largest effect in the right hemisphere areas.

We also observed a laterality effect in the difference between recalled and forgotten words, also known as the subsequent memory effect (Fig. 3C). Induced power was higher on trials with words that were subsequently recalled than on those that were forgotten. In the left hemisphere, this subsequent memory effect (SME) difference was increasing going from the most posterior to the anterior brain regions, reaching the peak magnitude in the anterior prefrontal cortex. The differences in the right hemisphere were relatively low and showed a peak memory effect in the MTL. Therefore, the SME confirms the hemispheric differences and overlap in the same brain regions that showed the largest laterality effect, including the inferior temporal cortex. This also highlights the importance of these higher association areas in memory function. This trend was not found in the right hemisphere, however, it is important to note that the highest right hemisphere SME was found in the MTL.

### 3.3. Slow, intermediate and fast activities show distinct spatiotemporal dynamics

In agreement with the previous studies (Burke, Long, et al. 2014; Burke et al. 2013; Kucewicz et al. 2014, 2019), the spectral power induced in this task followed a temporal sequence of brain regions, showing enhanced or suppressed power at specific times of word encoding (Fig. 4). We compared these sequences across the nine ROI in the six frequency bands with reference to the four phases of word encoding (PRE - before the word appears on the screen, EARLY - the first 800 ms of the word presentation, LATE - the second 800 ms the presentation, and POST - after the word disappears from the screen). First, we confirmed, while accounting for ROI, hemisphere, and frequency band, that the spectral power was significantly modulated by the phase of memory encoding (repeated measures ANOVA, F=729.45, p<001, df = 3). In each band, we found that the power alternates between a relative induction and a suppression, but the timing and the ROI sequences were different across the frequency spectrum. The slow theta, intermediate alpha/beta, and the fast gamma each showed a distinct pattern (Fig. 4A). The intermediate alpha and beta power was enhanced mainly in the pre- and the post-encoding phases. In contrast, the slow and the fast band power was induced in the early encoding phase in two waves of activation - first at word onset in the theta bands and then following the onset in the gamma frequencies. There was a significant effect of the frequency band (repeated measures ANOVA F=18.06, p<0.001, df=5), of the brain region (ANOVA F=15.57, p<0.001, df=37), and of the hemisphere location (ANOVA F=72.72, p<0.001, df=1) on the peak power latency. In the gamma bands, earlier latencies were observed in the visual compared to the prefrontal cortical areas (post-hoc Tukey-Kramer, p<0.05), which was not the case for the theta bands (Fig. 4B). The high gamma power peaks occurred significantly later than the theta peaks (post-hoc Tukey-Kramer, p<0.05) in the prefrontal cortical areas (Fig. 4C left) but not in the visual areas (Fig. 4C right), where the alpha/beta peaks followed later than the high gamma peaks. Therefore, the sequences of the induced power were different both between the slow and fast activities in the same cortical areas, and between the cortical areas in the same frequency bands.

**Figure 4.**
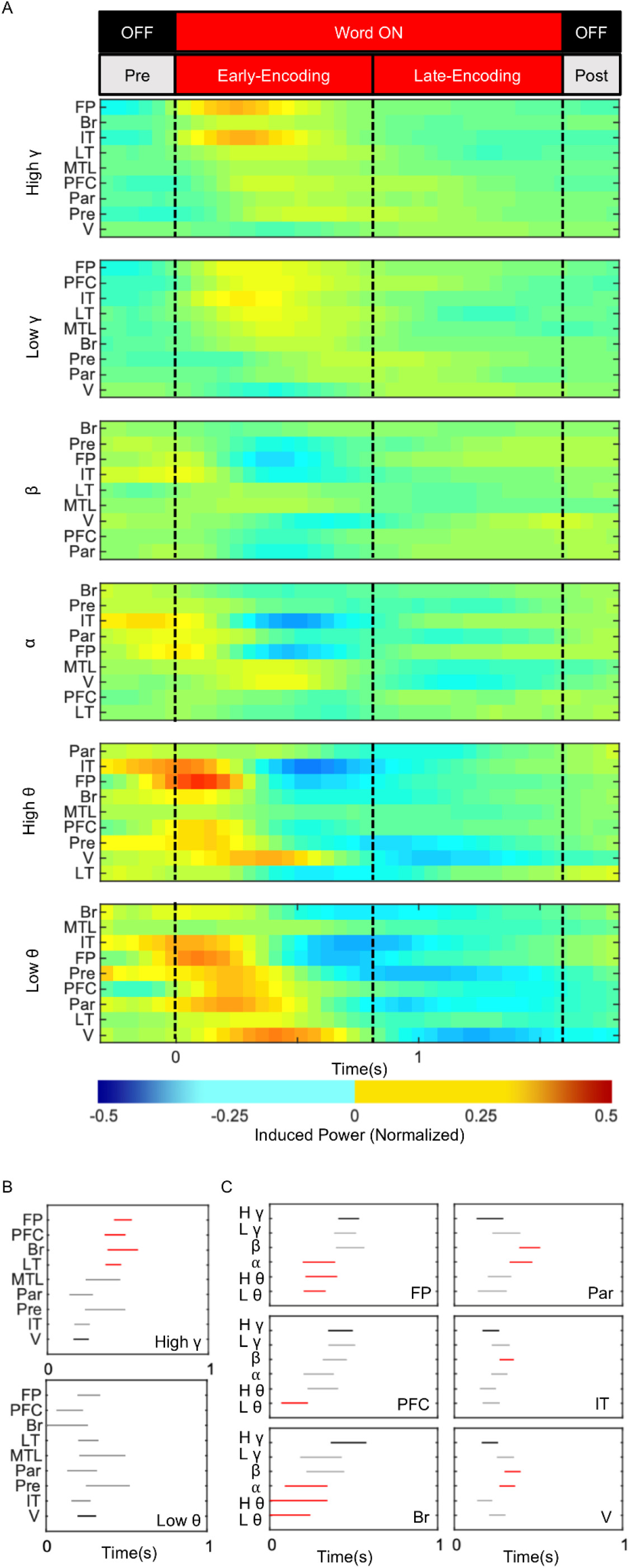
Induced slow, intermediate and fast activities are temporally arranged into distinct sequences of cortical brain regions. (A) Mean power changes estimated from correct recall trials of all active electrodes in a given brain region of the left hemisphere are ordered from the earliest to the latest peak latency (dashed lines separate different phases of memory encoding). Notice consistent sequences of activation in the theta, the alpha/beta, and the gamma frequency bands. (B) Summary of the mean latency ranges (post hoc ANOVA mean comparison) in the time of word presentation (0 corresponds to onset) shows significantly earlier gamma power induction in the most anterior FP areas (black) relative to the more posterior areas (red; p<0.05, Tukey-Kramer test), and an opposite pattern of the theta power induced earlier in the posterior V area relative to the more anterior brain regions (see Fig. 2 for labels). (C) Anterior and posterior brain regions show analogous patterns of power induced significantly earlier in the low frequencies of the prefrontal areas on the left, and significantly later in the alpha/beta frequencies of the visual areas on the right (red; p<0.05 Tukey-Kramer test) relative to the high gamma band (black).

Finally, to provide a complete picture of these spatiotemporal dynamics across the frequency spectrum we interpolated the induced power values from all active electrodes on an average brain surface. Spectral power was predominantly induced in the posterior visual and in the anterior prefrontal cortical areas at the encoding phases specific for the slow, intermediate and fast frequency activities (Fig. 5 top). Induced power was first observed in the theta activities of the visual areas before word onset and then right after the onset in the anterior prefrontal areas. This early posterior-to-anterior sequence of theta power induction was followed by a secondary induction of the fast gamma power in the same sequence later into the word encoding. These two sequences of induced power are propagating across the anatomical space in both hemispheres (Suppl. Video 1). There was no analogous propagation observed in the alpha or the beta bands. The alpha/beta power was induced in the same posterior and anterior areas but predominantly before and after word presentation, suggesting a more preparatory or inhibitory role in our proposed model of human memory encoding (Fig. 5 bottom). During preparation for encoding of the incoming words, the intermediate frequency oscillations would prevail and be followed by induction of the slow theta power across the brain at the time of word onset. The theta rhythms would in turn entrain the posterior and the anterior areas for final induction of the fast gamma activities during the time of word encoding, first in the visual and then progressively later in the higher-order prefrontal cortical areas (Burke, Long, et al. 2014; Kucewicz et al. 2019). The model is proposed to link the dynamics of particular spectral activities with the associated cognitive processes involved in anticipation, attention, and encoding of the remembered stimuli.

**Figure 5.**
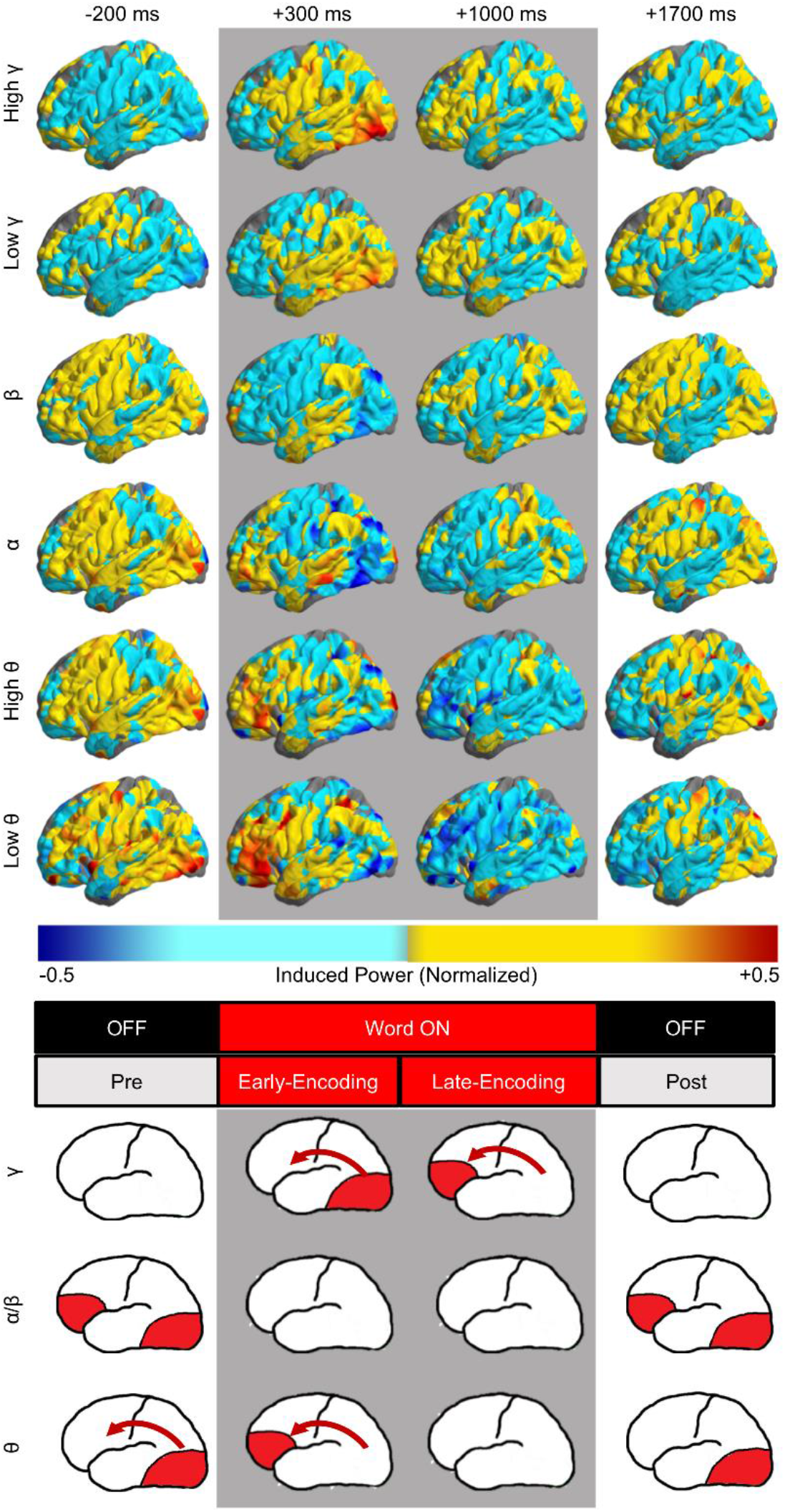
Anatomical spread of slow and fast activities propagates as distinct posterior-to-anterior waves of spectral power. (Top) Average brain surface plots summarize interpolated power from all active electrodes at selected four times points of the pre-, early, late, and post-word encoding periods (gray background indicates word presentation on the screen). Notice analogous patterns of the anatomical spread for the slow (theta), intermediate (alpha/beta), and fast (gamma) activities, especially in posterior visual and the anterior prefrontal areas. (Bottom) Model of independent activation of the slow and fast frequencies propagating from the posterior to the anterior cortical areas during word presentation, preceded and followed by the intermediate alpha and beta frequency band activation outside of the word encoding period. Red arrow indicates direction of propagation.

## 4. Discussion

Separate anatomical distributions of the slow, intermediate, and fast spectral activities in the human cortex is one of the main findings of this work. Because of the large number of participants and electrodes, implanted throughout all cortical regions, recordings from discrete and spatially distinct sites across 39 Brodmann areas was possible. With exception of the visual cortex, within any one area a majority of the electrode sites showed task-induced activity only in one or two bands of the frequency spectrum. At the same time, activities in all six bands were found among all sites within a given area, producing a “mosaic-like” pattern of theta, alpha, beta and gamma activities recorded at different sites. Each site could be viewed as one small tile of the mosaic with a specific color corresponding to a particular band as visualized in the Fig. 2 brain surface plots. Our results are in agreement with the spectral fingerprint view (Siegel, Donner, and Engel 2012) that processing in a given cortical site is characterized by specific neural activities, as previously demonstrated in the MEG and iEEG studies (Fellner et al. 2019; Keitel and Gross 2016). Various spectral activities were sparsely distributed in any one cortical region in more or less equal proportions, despite subtle differences, e.g., relatively more gamma band activities in the prefrontal cortex. This general pattern of heterogeneous, sparse distribution would explain the observation of a broadband ‘spectral tilt’ of power across all frequencies (Burke, Ramayya, and Kahana 2015; Voytek and Knight 2015; Kilner et al. 2005; Miller et al. 2014) when activities from multiple sites in a given brain area were averaged together. As a result, low frequency power (<30 Hz) is typically decreased and high frequency power (>30 Hz) is increased upon activation of a cortical area in a given task. This broadband tilt in the spectral power can be resolved into specific spectral fingerprints at individual electrode sites. One could go even further to ask how resolved this spectral distribution is and whether it can still be observed on the level of micro-electrode sites. Our study was limited to multiple macro-contacts that were grouped from different subjects - high-density recordings from multiple macro- and micro-electrodes implanted in individual subjects (Kucewicz, Berry, Worrell chapter in Lhatoo, Kahane, and Luders 2019) would be required to address these questions further and elucidate the neuronal activities underlying the spectral fingerprints.

The six frequency bands were similarly represented in the cortex, as quantified by the relative proportion of electrode sites recording theta, alpha/beta, and gamma activities. Even though cortical task activation was typically associated with spectral changes in the theta and gamma frequency bands (Miller et al. 2014; Greenberg et al. 2015; Osipova et al. 2006; Solomon et al. 2017; J. F. Burke et al. 2013; Kucewicz et al. 2014), we found the same or higher number of active electrode sites in the alpha and beta bands. Oscillations in these frequency bands were previously implicated with anticipatory or even inhibitory processes preceding task activations (Spitzer and Haegens 2017; Engel and Fries 2010). Compared with post-stimulus gamma activities related to sensory processing, alpha and beta activities were reported before stimulus onset and modulated by attention (van Ede, Szebényi, and Maris 2014; Bauer et al. 2014). Our results confirm the different timing of alpha and beta power induction, occurring mainly before and after the period of word presentation on the screen. These were observed in anatomical areas that were overlapping with the gamma and theta power induced during word presentation. Hence, the spatiotemporal pattern of spectral power confirmed distinct roles played by the theta, gamma, as well as the alpha and beta activities.

Theta frequency bands were also found to show a profile of spatiotemporal changes that was distinct from the gamma activities. Task-induced power in the high gamma band is known to arrange into a hierarchical sequence of brain regions activated first in the posterior sensory areas and then in the more anterior associational temporal and prefrontal cortex (Kucewicz et al. 2019, 2014). This posterior-to-anterior propagation of high gamma power inspired a two-stage model of memory encoding (Burke, Long, et al. 2014) to distinguish the early sensory and the late semantic phases of processing words. Theta power was also reported in the same areas preceding the gamma activation (Burke et al. 2013). Our recent analysis showed that this hierarchical activation from the sensory posterior areas to the anterior semantic systems propagates continuously along the ventral visual and semantic processing streams, ending in the ventrolateral prefrontal cortex of the Broca’s speech area and in the frontal pole (Kucewicz et al. 2019). Here, we observed a “wave” of induced power propagating across the cortex in the posterior-to-anterior sequence that was different for the theta and the gamma activities. First a theta wave was initiated at the time of stimulus onset in contrast to the gamma wave that started following the onset. In addition, the shortest latencies of the induced theta power peaks were localized in the Broca’s and prefrontal areas even before the visual cortex. The gamma power peaks, on the other hand, revealed the shortest latencies in the visual cortex and the longest in the Broca’s area, prefrontal cortex and the frontal pole. Hence, both the timing and the anatomical sequence of cortical areas was different for the power induced in the theta and the gamma bands. This suggests that the slow and fast activities are independently induced, playing different roles in stimulus processing. It remains an open question whether this propagation is in the form of actual travelling waves as described in the phase analysis of theta and alpha oscillations (Zhang et al. 2018), which confirmed the general posterior-to-anterior directionality of the local cortical waves. That study also pointed to very local discrete cortical locations of either slow theta, fast theta or alpha oscillations in all cortical areas. Existence of analogous local waves in the gamma activities on the same or a smaller scale is debatable, but most likely would be different from the slow activities.

We summarized this complex picture of the slow, intermediate, and fast neural activities in a simplified model of the relative spatiotemporal dynamics during memory encoding (see Fig. 5). Alpha and beta oscillations dominated in the initial pre-stimulus presentation phase potentially related to anticipatory and attentional processes in both the posterior sensory and the anterior association areas. Theta activities are then induced in these areas, we propose, in preparation for the expected stimulus processing around the time of presentation. Gamma activities are finally induced in response to the stimulus presentation in a sequence of the processing stream from the visual cortex to the prefrontal areas. This comprehensive view of the relative dynamics of these activities can be best captured using all dimensions of the spectral scale, anatomical space and time of stimulus processing (see Suppl. Video 1). Functional roles of these distinct spectral activations are not the subject of this study, although they are congruent with anticipatory, attentional, preparatory, and perceptual processes proposed in the literature. In general, the slower spectrum dominates in early phases of preparation and receiving stimulus information, whereas the late phase of encoding the presented information is characterized by gamma activities accompanied by suppressed power of the slower oscillations. Bursts of gamma power were previously described in humans and non-human primates in response to processing memory items (Kucewicz et al. 2014, 2017; Lundqvist et al. 2016, 2018). How these bursts of fast activities are related to the slower oscillations through, e.g. phase coupling, and understood in frames of various models like the theta-gamma memory buffer (Lisman and Jensen 2013) remains to be reconciled with our reported independent spatiotemporal dynamics. Our results pertain to the induced spectral power with little overlap between different frequencies and no evidence for amplitude-amplitude interactions. However, phase-phase or phase-amplitude interactions between the slow and the fast activities (Canolty et al. 2006) are still possible and could explain the capacity limit of the remembered items (Kamiński, Brzezicka, and Wróbel 2011), which was observed also in our study of free recall. Other bivariate measures of phase interactions, for instance, spectral coherence and network connectivity (Burke et al. 2013; Solomon et al. 2017), would also be required to provide a full picture of the neural dynamics engaged in memory formation.

Neural dynamics differed between the two hemispheres, revealing different profiles of spectral power, particularly in the prefrontal and the temporal cortex. The lateral prefrontal and inferotemporal cortices showed the greatest laterality effect and SME with more power in the left hemisphere, while the visual cortex had more power in the right hemisphere. In general, this hemispheric asymmetry was found to various degrees and in at least one frequency band in each brain region. Spectral fingerprints were thus specific to the hemisphere. One possible reason for this asymmetry is a differential engagement in the encoding and retrieval processes. According to the HERA (hemispheric encoding/retrieval asymmetry) model, we would expect brain activities during encoding to be stronger in the left hemisphere than in the right hemisphere and for the opposite to be true for retrieval (Buckner et al. 1996; Rugg et al. 1996; Fletcher, Shallice, and Dolan 1998; Fletcher et al. 1998; Grady et al. 1998). Therefore, it is not surprising that higher induced power was found in the left lateral prefrontal and inferior temporal cortex, especially in a verbal task with words. While a complimentary analysis of the induced power during retrieval of recalled words was beyond the scope of this study, others reported in the recall epoch of the same task more theta power in the right temporal and more gamma power in the left prefrontal and temporal cortex areas (Burke, Sharan, et al. 2014). Previous studies with different tasks (Tulving et al. 1994; Fletcher, Frith, and Rugg 1997) associated the right prefrontal cortex with episodic memory retrieval, and the left prefrontal cortex with encoding. Therefore, the preferential activation of the left prefrontal cortex during encoding in our task can be explained both within the HERA model and also by the verbalizability of the stimuli used (Golby 2001). The latter can be further studied with determination of language dominance for every subject.

Influence of the pathophysiology of epilepsy on the laterality effect or the estimates of spectral power in general can only be minimized but not completely eliminated in this patient population. Electrode channels that showed epileptiform activities in the seizure-generating areas were excluded from the analysis during the automated selection of active electrodes. Although occurrence of epileptiform activities was shown to be related with memory processing at distinct phases and anatomical locations (Matsumoto et al. 2013; Horak et al. 2017), it is highly unlikely that these pathological activities would consistently occur at specific phases of memory encoding and thus bias the power estimates used for selecting active electrodes. Hence, the laterality or the memory effects reported here are presumed to be only minimally affected.

Our results showed that it is not enough to generalize the neural activities to a given cortical area but a further distinction between the area in the left and the right hemisphere is critical. The cortical areas with the greatest laterality effect also showed a high memory effect. Spectral power induced on trials with words that were subsequently recalled was greater than on the ones with words that were forgotten in the left temporal and prefrontal cortex. This subsequent memory effect was gradually increasing in magnitude with successively more anterior cortical areas. Previous neuroimaging and electrophysiological studies found this memory effect to be the highest in the lateral prefrontal and the temporal cortex during similar tasks (Wagner et al. 1998; Long, Oztekin, and Badre 2010; Long, Burke, and Kahana 2014; Kucewicz et al. 2019; Kim 2011; J. F. Burke et al. 2013; Sederberg et al. 2003). Our recent investigation of the memory effect in the same task localized the highest magnitude in the left ventrolateral prefrontal cortex close to the Broca’s area, and in the left occipitotemporal cortex (Kucewicz et al. 2019). Altogether, the cortical areas identified with the greatest induced power and hemispheric laterality in the task overlap with localization of the memory effect.

Despite its known involvement in declarative memory function (Beason-Held et al. 1999); (Zola-Morgan, Squire, and Ramus 1994); (Parkinson, Murray, and Mishkin 1988); (Squire and Zola-Morgan 1991), MTL and the hippocampus had a relatively low number of active electrodes, magnitude of the induced spectral power, and laterality effect. One reason for this finding is a sparser distribution of hippocampal activities than in the neocortex, which could result in effectively less induced power recorded on any one macro-contact electrode. Sparse micro-contact neural activities would be averaged out by the surrounding non-active areas, compared to more topographically mapped cortical areas. Another reason is the task itself that may engage the MTL less than a more spatial or episodic memory task. Remembering the sequence of words on the list would be expected to increase hippocampal involvement in the task. A greater neocortical contribution to this task would explain our reports of enhanced performance in this task with stimulation in the lateral temporal cortex (Kucewicz, Berry, Miller, et al. 2018; Ezzyat et al. 2018). Stimulation in MTL was found to have an opposite effect (Kucewicz, Berry, Miller, et al. 2018; Jacobs et al. 2016), suggesting different roles for the two structures in this task.

Mapping neural activities that are critical for memory encoding guides the development and targeting of brain modulation, e.g. with the direct electrical stimulation, for therapeutic and research purposes. Out of the multitude of spectral activities that are induced at various times and anatomical locations it is necessary to identify prospective targets for modulation. Spectral power and its various derivatives, including the laterality or the memory effect, offer simple and accessible biomarkers for new brain-computer interface technologies. Such biomarkers can be rapidly computed to the brain regions, times and states for modulating brain activities underlying memory and other cognitive functions. For instance, they can be used to determine the neural activities correlated with memory enhancement (Kucewicz, Berry, Kremen, et al. 2018), to predict memory states for brain stimulation and rescue poor encoding trials (Ezzyat et al. 2017), or trigger modulation online in a closed-loop design for responsive electrical stimulation (Ezzyat et al. 2018). Hence, the spectral biomarkers can be useful for developing and optimizing the emerging technologies and new therapies for mapping and restoring memory functions.

## Acknowledgements

Cindy Nelson and Karla Crockett assisted in participant recruitment and data collection. This work would not be possible without the dedicated effort and participation of participants and their families. Open-access datasets were originally collected as part of a BRAIN Initiative project called Restoring Active Memory (RAM) funded by the Defense Advanced Research Project Agency (DARPA). This research was supported from the First Team grant of the Foundation for Polish Science co-financed by the European Union under the European Regional Development Fund.

## Supplementary Figures

**Supplementary Figure 1.**
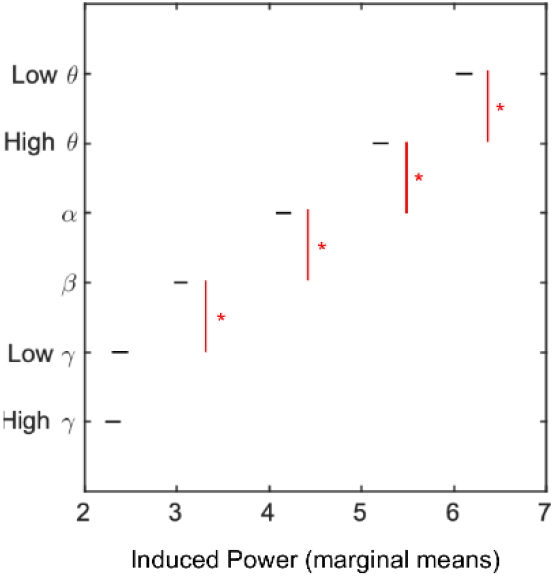
Post-hoc group analysis confirmed significantly smaller power with increasing frequencies of the bands, except between the low and high gamma bands. (Tukey-Kramer, p<0.05). Red lines with red asterisks indicate significance.

**Supplementary Video 1. An early posterior-to-anterior sequence of theta power induction is followed by a secondary induction of the fast gamma power in the same sequence later into the word encoding.** These two sequences of induced power are propagating across the anatomical space in both hemispheres. There was no analogous propagation in the alpha or beta bands. It is important to note that this video includes all word presentations, not just those that are later recalled, but a clear wave from posterior to anterior regions can still be seen in this video. Timing Bar running at top and bottom changes color to red to indicate word on. (L – Low, H – High)

